# Transmembrane coupling of protein condensates via membrane-mediated interactions: A simulation study

**DOI:** 10.64898/2026.08.14.744969

**Authors:** B. Ruşen Argun, Jeanne Stachowiak, Pengyu Ren

**Affiliations:** Department of Biomedical Engineering, University of Texas at Austin; Department of Chemical Engineering, University of Texas at Austin

## Abstract

Recent experiments show that protein condensates sitting on opposite surfaces of a flat lipid membrane move together and prefer to overlap, even though they cannot touch each other. This points to an indirect, membrane-mediated interaction. Two mechanisms could be responsible: a curvature-induced interaction, which is energetic in origin, and a fluctuation-induced interaction, which is entropic. Here we study both with coarse-grained molecular dynamics simulations, using Cooke’s implicit-solvent lipid model together with a generic bead-spring polymer model for the condensate. We compute the potential of mean force between two condensates across the membrane. For condensates of the same size, full overlap is unfavorable, and the pair instead settles into a partially overlapping state that bends the membrane into an S-like shape. When the two condensates differ strongly in size, full overlap becomes favorable. We explain this with a simple geometric picture. The condensate wets the membrane as a thin film and imposes curvature only along its rim, while membrane tension flattens the membrane under its interior. The resulting ring of curvature can trap a smaller condensate on the opposite side. We also compare the bending undulations and the effective bending modulus of a bare membrane, a membrane with one condensate, and a membrane with condensates on both sides. A wetting condensate suppresses the undulation modes and stiffens the membrane, but whether this makes overlap entropically favorable remains inconclusive. Our results indicate that the coupling is driven mainly by curvature, and that it depends on the wetting mechanism and on the membrane tension.

## I. INTRODUCTION

Liquid-like condensates of intrinsically disordered proteins (IDPs) can form in the nucleus and the cytoplasm, where they carry out a range of biological functions [1–3]. These condensates can adhere to and reshape membranes, mediating other membrane interactions [4–8].

Condensates have been observed to couple across membranes, although the mechanism responsible for this coupling remains poorly understood. Specifically, Lee *et al*. [9, 10] showed that protein condensates on opposite sides of a flat lipid bilayer diffuse together along the membrane surface, despite being unable to interact directly. This coordinated motion suggests an indirect, membrane-mediated interaction that generates an effective attraction between the condensates. The physics behind this phenomenon is worth investigating, as it could also be exploited by living cells as an alternative mode of communication between intra- and extracellular regions, one that requires neither a pore, nor a signaling protein, nor material transport.

Although membrane-mediated interactions between rigid membrane inclusions, such as membrane-embedded proteins, have been extensively studied in both experiments and simulations [11–13], the corresponding interactions between condensates remain less well understood. The literature suggests two main membrane-mediated mechanisms that could drive this coupling: curvature-induced and fluctuation-induced interactions [14]. Curvature-induced interactions are energetic in origin: the membrane favors configurations that reduce its bending energy. Fluctuation-induced interactions are entropic in origin: at finite temperature, membrane fluctuations favor macrostates with greater entropy. While these mechanisms have primarily been studied for membrane-embedded proteins, much of the underlying physics should also apply to condensates wetting a membrane, even though this represents a distinct physical setting.

The fluctuation-induced interaction is attractive between inclusions on a lipid membrane [14]. Rigid inclusions suppress the membrane’s bending modes, so bringing them close together lets the membrane recover the otherwise suppressed modes, increasing its entropy; the effect is analogous to the depletion force between colloidal particles and to the Casimir effect [15]. A similar entropic mechanism has been suggested to couple nanodomains across opposing leaflets of a bilayer [16]. Haataja [17] explained the co-localization of membrane domains on opposite leaflets by arguing that rigid lipid domains suppress bending undulations, so placing two such domains at the same lateral position minimizes the total suppression of membrane modes and maximizes entropy. They used Helfrich theory and mode-counting arguments to estimate the resulting coupling. Related behavior has also been observed in particle-based simulations: Pezeshkian *et al*. [18] modeled toxins as rigid nanoparticles adhering to a bilayer and, using coarse-grained DPD, argued that suppression of thermal membrane fluctuations is the primary driver of their clustering; Madsen *et al*. [19] used a five-bead-per-lipid model and concluded that membrane entropy can drive the clustering of transmembrane proteins; and Jafarinia *et al*. [20] showed that a fluctuation-induced interaction aggregates transmembrane proteins in a rigidity-dependent manner. Although fluctuation-induced interactions are often argued to be negligible in most biological settings, at least for trans-membrane inclusions [21], they may be more relevant for the micron-sized condensates seen in the experiments.

In the curvature-induced picture, wetting by a condensate creates a local force imbalance where the condensate, membrane, and surrounding solvent meet. The membrane can relax this imbalance by bending, analogous to the curvature imposed by rigid membrane inclusions [8, 22, 23]. Van Der Wel *et al*. [24] showed, through experiments and dynamically triangulated Monte Carlo simulations, an effective attraction of about 3 *k*_*B*_*T* between pairs of micron-sized colloidal particles adhering to and bending an elastic membrane, and concluded that the interaction is independent of particle size. Theoretical studies find curvature-induced interactions of comparable magnitude, and related work has modeled deformable particles and vesicles adhering to membranes [25, 26].

The net interaction between two membrane-deforming objects can be either repulsive or attractive, depending on the sign and magnitude of the curvature they impose on the membrane [27–29]. Therefore, when attraction is observed, it is not immediately clear whether the coupling is driven by curvature, fluctuations, or both. Sadeghi [30] used coarsegrained simulations of curvature-inducing peripheral proteins and showed that the entropic attraction can dominate the energetic curvature repulsion and drive aggregation. Cleanly separating the two contributions in a simulation remains challenging, however, because they are strongly coupled [28]. How, then, do these two mechanisms contribute to the transmembrane coupling of condensates?

In this work we address this question with a highly coarsegrained model of the membrane–condensate system. We perform potential-of-mean-force calculations to quantify the strength of the coupling under different conditions, such as membrane tension and condensate size asymmetry. We also quantify the bending undulations of a bare membrane, a membrane with a condensate on one side, and a membrane with condensates on both sides, in order to assess whether a fluctuation-induced interaction can play a role and whether a condensate increases the membrane’s effective bending modulus. In the discussion, we connect our findings with previous theoretical expressions for the two mechanisms. Throughout, we combine coarse-grained and continuum descriptions, as multiscale analysis is needed to span the wide range of length and time scales involved in protein–membrane interactions [31].

## II. METHODS

### Coarse-Grained Model for Membrane and Condensates

Lipid bilayer membranes can be modeled at a wide range of resolutions, from atomistic to continuum [32–34]. Energy-minimizing elastic continuum models can capture the curvature-induced effects that may contribute to coupling, but fluctuation-induced interactions require a model that also includes thermal noise. Protein condensates can likewise be modeled at atomistic [35], coarse-grained [36], and continuum [37] resolutions. Here, our goal is to investigate membrane-mediated interactions between two liquid-like condensates composed of protein chains. This objective imposes lower bounds on the relevant length and time scales, since membrane-mediated interactions, particularly in free-energy calculations, require long equilibration times. Explicit-solvent models are therefore too expensive for our purposes, even at coarse-grained resolutions such as Martini, despite being relatively straightforward to implement.

We instead use a highly coarse-grained implicit-solvent membrane model with several beads per lipid, which provides a practical balance between physical resolution and accessible length and time scales. For a 100nm × 100nm membrane patch, implicit-solvent models typically require roughly an order of magnitude fewer particles than explicit-solvent representations. Specifically, we use Cooke’s model [38, 39], a three-beads-per-lipid model with generic Lennard-Jones-like nonbonded interactions. Because Cooke’s model is chemistry-agnostic, a natural counterpart for the protein condensate is a bead-spring coarse-grained polymer model, which can be described using similarly generic nonbonded interactions.

Here we describe the details of the model. All quantities are reported in standard Lennard-Jones (LJ) reduced units, with the lipid tail bead defining the unit of length *σ* and the characteristic non-bonded interaction strength defining the unit of energy *ε*. Temperatures are reported in units of *ε/k*_B_, and times in units of 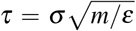, where *m* is the bead mass (taken to be unity for all species). Simulations were carried out in a rectangular box of dimensions *L*_*x*_ = *L*_*y*_ = *L*_*xy*_, with periodic boundary conditions in all three directions. A box of *L*_*xy*_ = 200.0*σ* is used for unbiased runs and umbrella sampling unless specified otherwise. The bilayer normal is aligned with the *z* axis. The bilayer is represented by the implicit-solvent, three-bead lipid model of Cooke and co-workers [38, 39], in which each lipid is composed of one head bead (type H) and two tail beads (type T) connected linearly, as depicted in Fig. 1 (a). Consecutive beads along a lipid are linked by a finitely extensible nonlinear elastic potential with a Weeks–Chandler–Andersen (FENE-WCA) repulsive core,

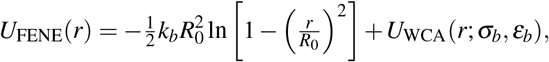

with *k*_*b*_ = 30*ε/σ* ^2^, *R*_0_ = 1.5*σ*, and *ε*_*b*_ = 1*ε*. The bond reference diameter *σ*_*b*_ is set to 0.95*σ* for head–tail bonds and to 1.0*σ* for tail–tail bonds. To enforce an extended lipid configuration we impose a harmonic bond potential between the head and end tail beads,

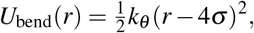

with bending stiffness *k*_*θ*_ = 10*ε/σ* ^2^. Non-bonded interactions among lipid beads consist of a short-ranged repulsion plus, exclusively for tail–tail pairs, an additional attractive tail that mediates bilayer self-assembly in the absence of explicit solvent. The repulsion is taken as a shifted Lennard-Jones (WCA) potential truncated at *r*_*c*_ = 2^1*/*6^*σ*_*i j*_, with *σ*_HH_ = *σ*_HT_ = 0.95*σ* and *σ*_TT_ = 1.0*σ* . For tail–tail pairs we superimpose the cosine attractive well of the Cooke model,

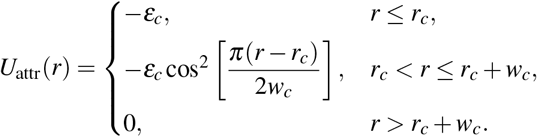

with *ε*_*c*_ = 1*ε, r*_*c*_ = 2^1*/*6^*σ*, and tunable range *w*_*c*_ = 1.6*σ* . This combination of WCA repulsion and cosine attraction reproduces the canonical Cooke parametrization that yields a fluid, self-assembled bilayer at the temperatures considered here. The bilayer is initialized as a flat hexagonal lattice with an areal lipid density of *ρ*_*ℓ*_ ≈0.85*σ* ^−2^ per leaflet.

**FIG. 1:**
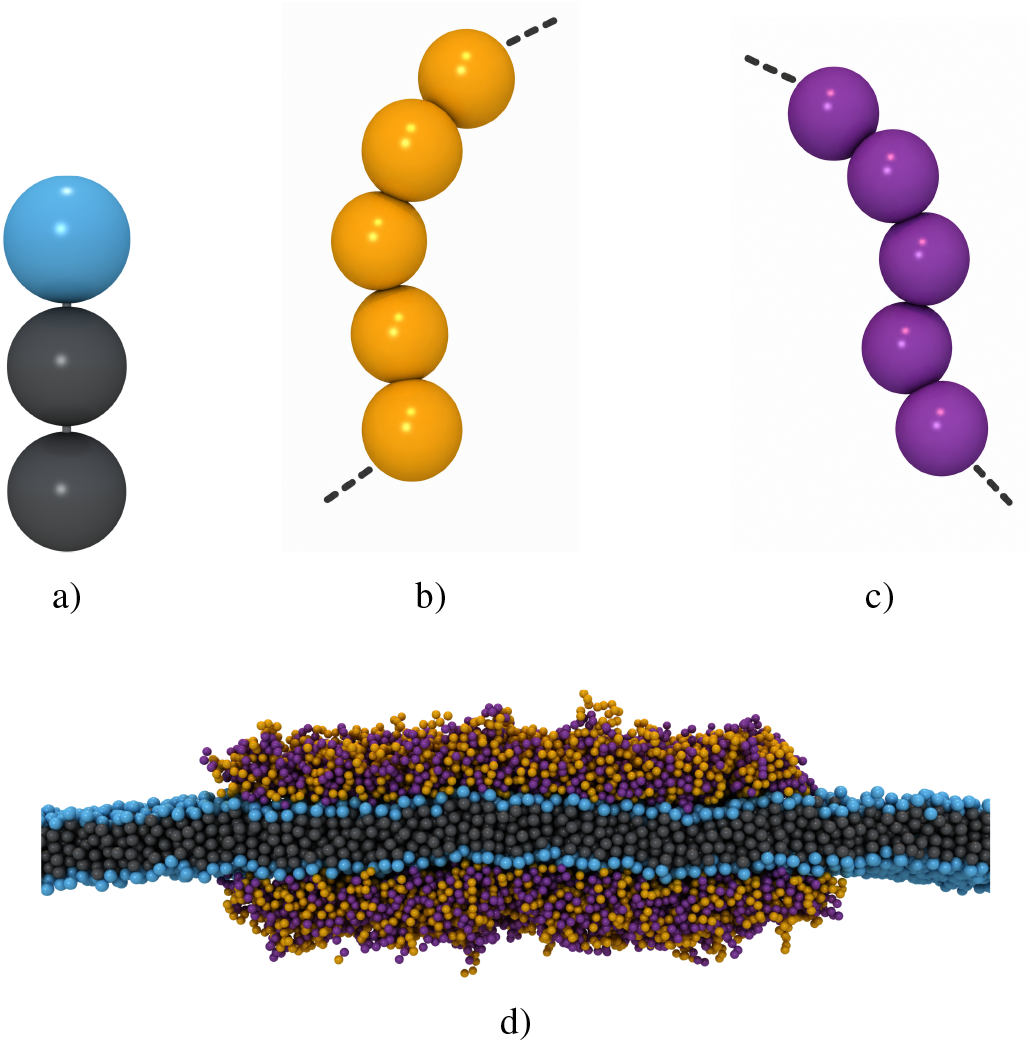
(a) Cooke’s lipid, one head bead and two tail beads. (b) Polymer chain with type A beads. (c) Polymer chain with type B beads. (d) Two condensate droplets wetting the membrane, cross-section view.

The intrinsically disordered protein molecules are modeled as bead-spring polymer chains [40–42]. Two polymer condensates, one grafted to each leaflet, are shown in Fig. 1 (d). Each polymer chain has 30 beads, unless specified otherwise, and the number of polymer chains in each condensate is denoted as *N*_*chain*_. Each chain is a homopolymer of one of two species *A* and *B*, shown in Fig. 1 (b) and (c) respectively [43]. Each condensate contains equal numbers of *A* and *B* chains. Consecutive monomers are connected by a stiff harmonic bond,

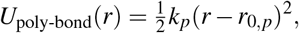

where *k*_*p*_ = 100*ε/σ* ^2^, and equilibrium length *r*_0,*p*_ = 0.7*σ* . Non-bonded interactions between polymer monomers are described by LJ potentials with diameter *σ*_*p*_. By matching the bilayer thickness to real lipid membranes, the effective size of the Cooke beads, *σ*, is approximately 0.9 nm [44]. A reasonable Kuhn length for IDP chains is ≈1.0 nm [45]. Since a bead-spring polymer with excluded-volume interactions has a Kuhn length of approximately 1.7 bead diameters, we set *σ*_*p*_ = 0.70*σ* .

Pairs of identical species (*A*–*A* and *B*–*B*) interact through a purely repulsive WCA potential (*ε*_*AA*_ = *ε*_*BB*_ = 1*ε*, cutoff 2^1*/*6^*σ*_*p*_), whereas unlike pairs (*A*–*B*) are treated with the full attractive 12–6 LJ potential,

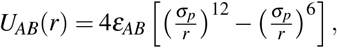

shifted to vanish at *r*_*c,AB*_ = 2.5*σ*_*p*_. At *k*_B_*T* = 1*ε* the cross-attraction *ε*_*AB*_ = 0.9*ε* lies just above the dense–dilute coexistence window (observed at *ε*_*AB*_ = 0.85*ε*), so each chain set condenses fully into a single liquid-like droplet on its leaflet. At this cohesion strength, we still observe significant shape fluctuations and deviations from a disk-like shape for the condensates, with polymers diffusing within them, indicating a liquid-like structure for the droplets.

Each polymer chain is anchored to the bilayer by a covalent tether between the chain’s terminal monomer and a selected head bead of the proximal leaflet in order to mimic experimental wetting conditions. Tether bonds are harmonic with the same stiffness as the polymer backbone, *k*_*t*_ = 100*ε/σ* ^2^, and equilibrium length *r*_0,*t*_ = 1.0*σ* .

Cross-species interactions between polymer beads and lipid beads are also of LJ type, with diameter *σ*_*p*H_ = 0.85*σ* for polymer–head pairs and *σ*_*p*T_ = 1.0*σ* for polymer–tail pairs.

Polymer–tail interactions are kept purely repulsive (WCA, cutoff 2^1*/*6^*σ*_*p*T_), so that polymer beads cannot insert into the hydrophobic core. Polymer–head interactions carry a tunable attractive component of strength *ε*_*p*H_ = 0.2*ε*, truncated and shifted at 2.5*σ*_*p*H_, which controls the adhesion of each condensate to its underlying leaflet. Setting *ε*_*p*H_ →0 recovers a non-adhesive (WCA) polymer–head interaction.

The equations of motion were integrated with a velocity-Verlet scheme using a time step Δ*t* = 5 × 10^−3^*τ*. Temperature was maintained with a stochastic velocity-rescaling thermostat [46] with relaxation time *τ*_*T*_ = 20Δ*t* = 0.1*τ*. The simulation cell was coupled to a Martyna–Tobias–Klein barostat [47] with characteristic time *τ*_*P*_ = 1000Δ*t* = 5*τ*, configured such that *L*_*x*_ and *L*_*y*_ are coupled and rescale together while *L*_*z*_ is held fixed, thereby imposing a prescribed mechanical surface tension Σ on the bilayer. Specifically, the lateral target stress was set to *P*_*lat*_ = (*P*_*xx*_ + *P*_*yy*_)*/*2 = *P*_*z*_ − Σ*/L*_*z*_, with normal target pressure *P*_*z*_ = 0, thereby imposing the desired bilayer tension through Σ = − *P*_*lat*_*L*_*z*_. HOOMD-blue v5.4.0 [48] with tree neighbor-list [49, 50] implementation was used to perform the simulations.

To determine whether coupling is thermodynamically favored, and by how much, we calculated the potential of mean force (PMF) between condensates on opposite sides of the membrane as a function of the distance between their centroids in the *xy* plane, *r*_*xy*_, as shown in Fig. 2. Because condensates diffuse and relax slowly on the membrane surface and interact strongly, enhanced sampling is needed to converge the PMF, even for small condensates. We use the classical combination of umbrella sampling and weighted histogram analysis. The reaction coordinate is sampled in consecutive windows of *r*_*xy*_, with a harmonic bias potential [51]. The individual histograms are then merged into a smooth PMF profile by the weighted histogram analysis method [52, 53], with the *k*_*B*_*T* ln *r*_*xy*_ Jacobian of the 2D radial coordinate added. Implementation details on equilibration, convergence, and error estimation are given in the SI.

**FIG. 2:**
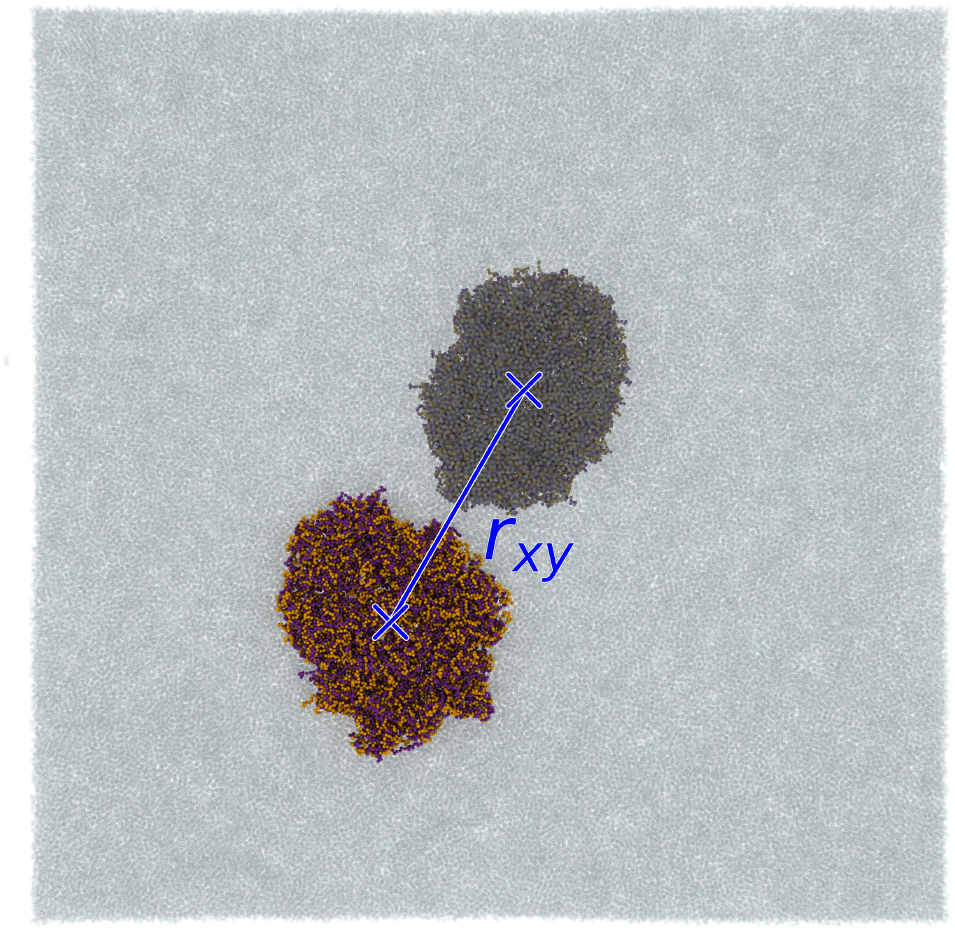
The reaction coordinate *r*_*xy*_ used for PMF calculations. The condensate on the lower left is above the membrane, the other condensate with dimmer colors is below the semi-transparent flat lipid membrane.

## III. RESULTS AND DISCUSSION

### Same-sized condensates

We first attempted to reproduce the coupling phenomenon observed in the experiments. When the two condensates were initialized *r*_*xy*_ = 0 on the flat membrane, we observed their coupled diffusion. To quantify the thermodynamic strength of this coupling, we computed the potential of mean force. In contrast to these unbiased trajectories, *F*_PMF_(*r*_*xy*_) revealed a strong, repulsive interaction between the two condensates (Fig. 3) at very short range. Longer MD simulations resolved this apparent contradiction: the condensates eventually drift apart to a finite offset even when started from the same *xy* location.

**FIG. 3:**
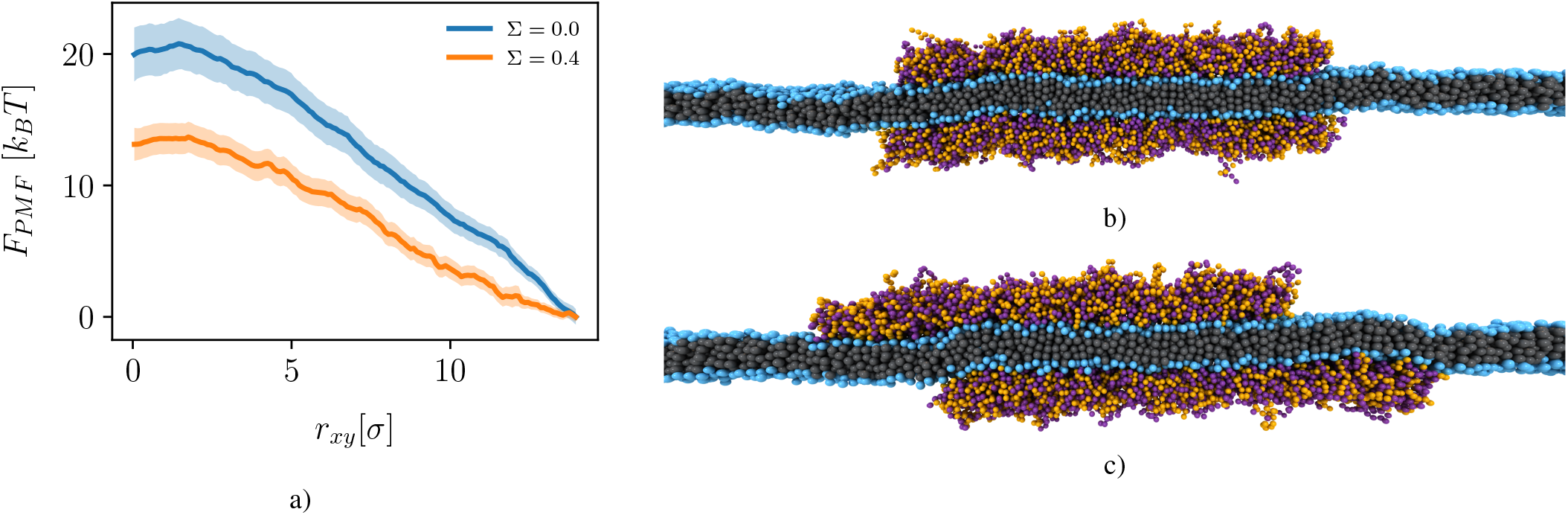
(a) *F*_PMF_(*r*_*xy*_) between two condensates with *N*_*chain*_ = 800 at lateral tensions Σ = 0.0 *k*_*B*_*T/σ* ^2^ and 0.4 *k*_*B*_*T/σ* ^2^. Shaded bands denote the statistical uncertainty. The zero-reference is taken as the minimum value of each *F*_PMF_(*r*_*xy*_) (b,c) Cross-section of configurations sampled at *r*_*xy*_ = 0*σ* (b) and 15*σ* (c).

We believe that the coupled (while being fully overlapped) diffusion seen in the MD trajectories, is largely due to friction between the two membrane leaflets. Because most lipids under a condensate on a given leaflet are either tethered directly to a polymer or neighbor a tethered lipid, the collective diffusion of one condensate drags the majority of that leaflet’s lipids along with it. The inter-leaflet friction drags the opposing leaflet and the other condensate. Even with our highly coarse-grained lipid model, this inter-leaflet friction makes equilibration slow.

Figure 3 demonstrated that fully overlapping configurations were not thermodynamically favored in our system. Because of the computational cost, we sampled *F*_PMF_ only over *r*_*xy*_ ∈ [0*σ*, 15.0*σ*], well below the approximate condensate radius (*R* = 30*σ* for *N*_*chain*_ = 800), two condensates still partially overlap at *r*_*xy*_ = 15.0*σ* as can be seen in Fig. 3(c). To probe for a local minimum beyond this range, we ran unbiased MD from several initial separations in [15.0*σ*, 30.0*σ*]. All trajectories converged to *r*_*xy*_ ≈23.0*σ*, (Fig. 4). For smaller condensates (*N*_*chain*_ = 500), a longer-range PMF shows this local minimum to be around 6 *k*_*B*_*T* deep (Fig. 5). Membrane-mediated interactions can thus couple the two condensates, though with one subtle distinction between simulation and experiment: whereas the experimental images [9] suggest a perfect coupling in which the projected condensates completely overlap within experimental resolution, the coupling we observe is offset, with the projected areas remaining very close but not fully overlapping (Fig. 4).

**FIG. 4:**
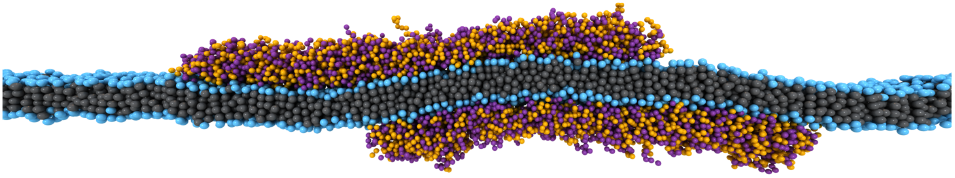
Equilibrium configuration (*r*_*xy*_ ≈23.0*σ*) of two same-sized condensates (*N*_*chain*_ = 800, Σ = 0.4*k*_*B*_*T/σ* ^2^): a cross-section of the system snapshot through the centers of the two condensates.

**FIG. 5:**
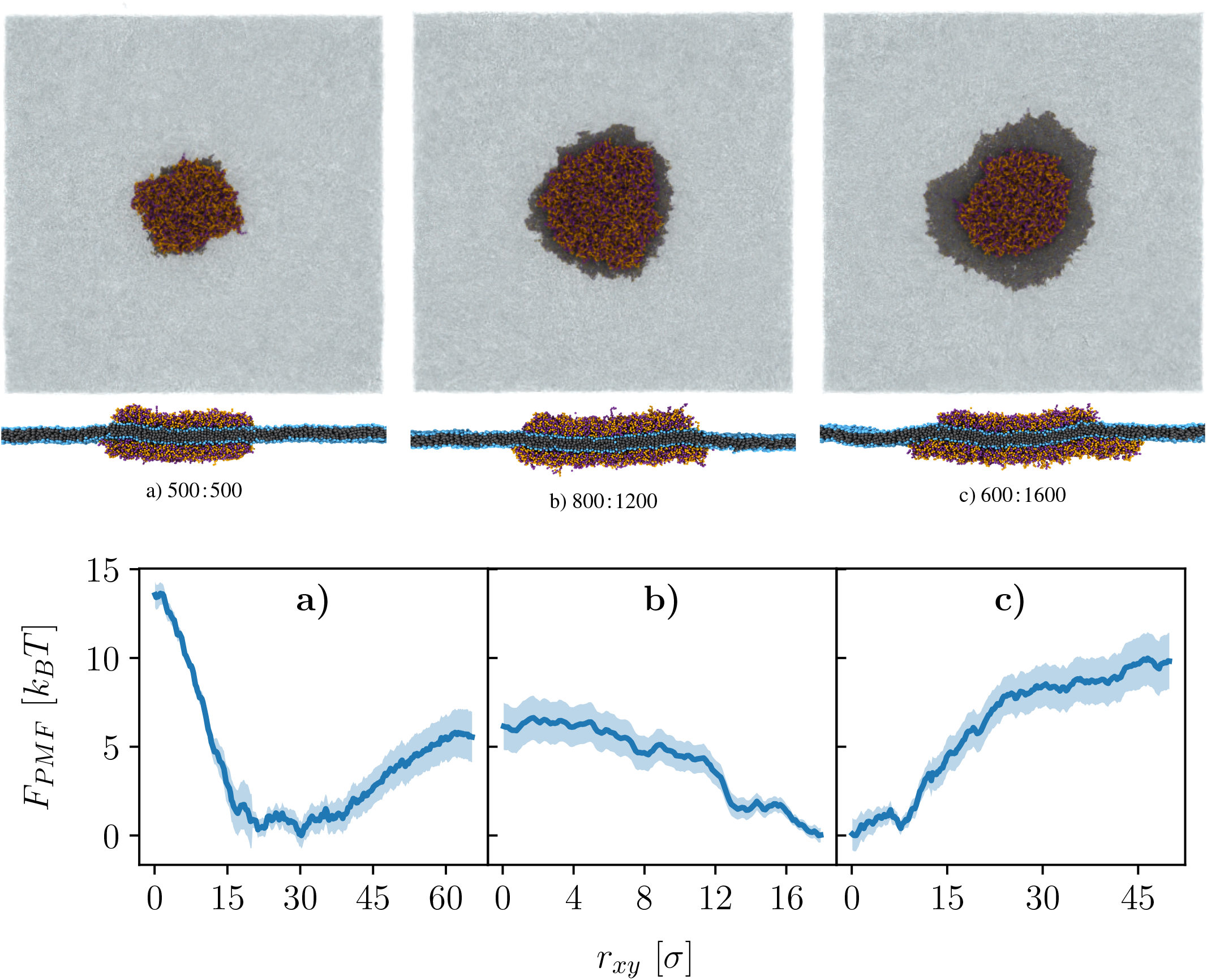
Effect of condensate size asymmetry on the membrane-mediated interaction. Columns correspond to (a) two equal condensates (500 : 500), (b) a small size mismatch (800 : 1200), and (c) a large size mismatch (600 : 1600), where the ratios denote the number of polymer chains in the upper and lower condensate, 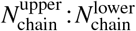 . *Top row:* top-down views of the condensate pair. *Middle row:* cross-sectional (side) views showing the membrane deformation. *Bottom row:* the corresponding potential of mean force *F*_PMF_(*r*_*xy*_) for cases (a)–(c), left to right; shaded bands denote the statistical uncertainty. Note the differing horizontal ranges across the three PMF panels.

The shape of the membrane cross-section in Fig. 4 hints that this coupling is driven by the membrane minimizing its bending energy in response to the opposing curvatures imposed by the two condensates. Although our configurations differ from the experimental ones, similar behavior has been reported previously: cooperative wrapping of curvature-imposing particles placed on opposite sides of a flat membrane yields results comparable to ours [54]. Using the Martini model, Mondal and Cui [55] simulated polyelectrolyte coacervates with lipid bilayers, and although membrane-mediated interactions were not their focus, their coacervates curve the membrane in a similar way when on opposite sides (Fig. 7 therein). On a much larger length scale, the continuum phase-field model of Mokbel *et al*. [56] likewise shows opposite-side condensates sitting next to one another while bending the membrane into an S-shape resembling Fig. 4, albeit with more pronounced curvature relative to the condensate size (their Fig. 4.6).

**FIG. 6:**
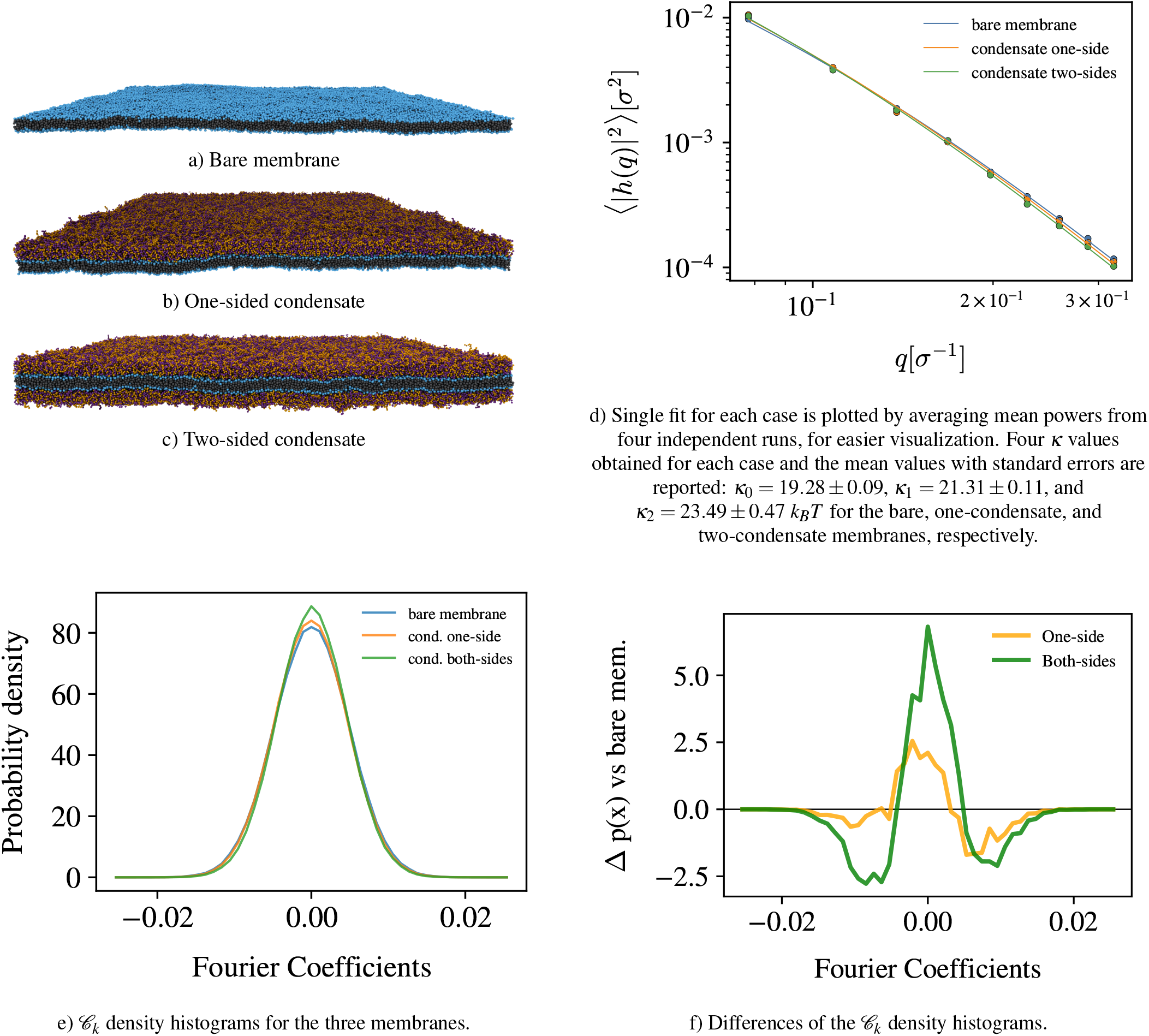
Membrane snapshots and bending undulation comparison. 4 independent runs for each case at the same tension, Σ = 0.4 *k*_*B*_*T/σ* ^2^, are performed to sample membrane height profiles. *L*_*xy*_ = 194.6 ± 0.2*σ*, 193.8 ± 0.2*σ*, 193.1 ± 0.2*σ* for bare, one-sided and two-sided cases, respectively. (a–c) Snapshots under three conditions; (d) *q* values within [0.08, 0.32]*σ* ^−1^ range are used for the fit. (e) *C*_*k*_ density histograms, which contain the Fourier coefficients of the transformed height fields *ĥ*^(*t*)^(**q**) = *A*^(*t*)^(**q**) + *i B*^(*t*)^(**q**) sampled within *q* = [0.4, 0.42]*σ* ^−1^ (f) The difference between *C*_*k*_ density histograms of 3 cases, bare membrane taken as the reference, same data as (e).

In principle, the umbrella-sampling/WHAM PMF can be decomposed into energetic and entropic contributions, which would be informative here since the membrane-mediated interactions are both energetic and entropic. In practice this is usually impractical, as the energy of the system is highly noisy and would require orders of magnitude more sampling to converge. That said, the continuum model of Mokbel *et al*. [56] neglects thermal fluctuations, and hence fluctuation-induced interactions, yet reproduces the same geometry, supporting our intuition that the observed coupling is primarily curvature-driven.

The suspended membranes in experiments are under non-negligible tension [9]. Comparing *F*_PMF_ with and without lateral tension (Fig. 3), we find that tension weakens the membrane-mediated interaction within *r*_*xy*_ = [0.0*σ*, 15.0*σ*] range. This is consistent with the general consensus that curvature-induced interactions are screened or weakened by membrane tension.

While our results agree with previous simulations, the mismatch with experiments and the competition between curvature- and fluctuation-induced interactions remain to be resolved, a challenging task as noted in earlier studies [28]. Many parameters influence both types of interaction, including the size of the condensates, their relative size, the membrane tension, and the curvature each condensate imposes on the membrane.

### Different-sized condensates

Looking carefully at experimental images ([9], Fig. 3 therein) we notice two things: condensates on one side of the membrane are usually larger than those on the other side, i.e. there is a size asymmetry, and the smaller condensates, while *fully* overlapping with the other, prefer to sit near the boundary of the larger condensate with which they are coupled. The reason for the size asymmetry is experimental; proteins are released to one side and it takes time for them to diffuse to the other side of the membrane which delays the condensation [9]. The tendency of the smaller condensate to sit at the periphery of the larger one points towards a more curvature-induced coupling mechanism as this is difficult to rationalize with fluctuation-induced mechanisms. For a purely entropic coupling we would expect all fully overlapping configurations to be degenerate. To see whether size asymmetry of condensates has any impact on the effective membrane-mediated interactions we repeated the PMF calculation with condensates of different sizes, at Σ = 0.4*k*_*B*_*T/σ* ^2^. The results are shown in Fig. 5, on the left panel we have same-sized condensates 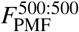, in the middle, condensates with small asymmetry 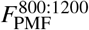, and on the right condensates with large size asymmetry *F*^600:1600^, where the integers on the superscript denote the number of polymer chains of condensates, 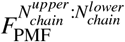.

We can see that the size difference between two condensates can significantly alter the effective interactions. While for same-size condensates there is an effective repulsion larger than 10 *k*_*B*_*T* that disfavors the completely overlapping configurations, this repulsion is reduced to ∼ 6*k*_*B*_*T* for condensates that contain 800 and 1200 polymer chains. The very short range repulsion is less steep as shown in Fig. 5 (a) vs (b). By increasing the size mismatch, we obtain a PMF that shows a depth of 10*k*_*B*_*T* for fully overlapping configurations, Fig. 5 (c).

How can a size difference reverse the effective interaction from repulsion to attraction? The visual inspection of cross-sectional snapshots suggests a simple geometric answer: if the small condensate is small enough to fit inside the curved membrane region created at the rim of the large condensate, the overlapping configuration becomes favorable. This is impossible for equal-sized condensates. Favorable curvature can only be achieved when they are partially separated, as in Fig. 4. In the experiments, however, even condensates that look similar in size are coupled, which raises the question of how much asymmetry is actually needed. Answering this requires clarifying how the condensate curves the membrane, and how this depends on the membrane tension (Σ), the condensate–solvent interfacial tension (Σ_*cs*_), the bending modulus (*κ*), the condensate radius (*R*), and the wetting mechanism.

In our model, the polymer–head attraction is weak (*ε*_*pH*_ = 0.2*ε*), so wetting is driven mainly by the tethering rather than by physical adhesion. Every chain is anchored, so the condensate is effectively a grafted layer, imitating the experimental system. The condensate is therefore not a three-dimensional droplet but an almost two-dimensional disk that spreads over the membrane surface, as the cross-sections show. This geometry lets us neglect the Laplace pressure exerted by the condensate on the membrane. The Laplace pressure is Δ*P* = Σ_*cs*_(1*/R*_1_ + 1*/R*_2_), where *R*_1_ and *R*_2_ are the principal radii of curvature of the condensate-solvent interface. For the interior of the condensate this interface is essentially flat, so Δ*P* vanishes and the bulk condensate exerts no distributed bending load on the membrane. All of the action is where the interface bends sharply, along the condensate rim: the curvature is imposed by a line force, not by a bulk pressure. Its magnitude is set by the competition between Σ_*cs*_ and *κ*, giving a curvature radius of order 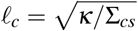 [4]. How far this deformation reaches into the surrounding membrane is set by a second length scale, a curvature screening length (or *bending boundary layer*) 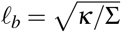, which follows from the competition between tension and bending.

For a tensionless membrane *ℓ*_*b*_ diverges, so the curvature imposed at the rim spreads over the whole membrane, and the Helfrich Hamiltonian is scale invariant. To illustrate this we compare two cross-sections (Fig. S3) in which the box size (*L*_*xy*_ = 150*σ*, 450*σ*) and the condensate size (*N*_*chain*_ = 444, 4000; with 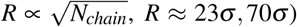 are scaled by the same factor. The membrane is bent everywhere, including the interior far from the rim, and the two membrane shapes look approximately like scaled versions of each other. Scale invariance means that for tensionless systems, small simulations (tens to hundreds of nm) are relatively more informative about the micron-scale experimental behavior. Applying tension makes *ℓ*_*b*_ finite and breaks this invariance, and the membrane shape relative to the condensate now differs between the two sizes (Fig. S4). For the Cooke membrane, *κ* ≈ 20*k*_*B*_*T* (estimated with eq. (10) in the next section) and if Σ = 0.4*k*_*B*_*T/σ* ^2^, we have *ℓ*_*b*_ ≈7*σ* . For the small condensate (*N*_*chain*_ = 444, *R* = 23*σ*) we have *R/ℓ*_*b*_ ≈3.3: the rim curvature is not damped before it reaches the interior and the membrane is bent as a whole under the condensate. For the large condensate (*N*_*chain*_ = 4000, *R* = 70*σ*), *R/ℓ*_*b*_ ≈10 and the tension has enough room to flatten the membrane under the interior, apart from thermal fluctuations. This directly controls how much asymmetry is needed for coupling: In the *R/ℓ*_*b*_ ≫ 10 regime even a small size asymmetry between the condensates would be sufficient for favorable bending with fully overlapping configurations. In the smaller *R/ℓ*_*b*_ ≲ 10 regime, which we probe with our simulations, larger size asymmetries are needed.

Turning back to the experiments, the bending modulus of biological membranes varies little and stays within 10 − 30*k*_*B*_*T*, while the membrane tension can vary by orders of magnitude. For suspended membranes used in the experiments [9] a typical value is Σ ≈1 mN*/*m [57], giving *ℓ*_*b*_ ≈ 10 nm, close to the simulation value of *ℓ*_*b*_ ≈ 7*σ* ≈ 6.3 nm. With micron-sized condensates *R* = 1*µm* this gives *R/ℓ*_*b*_ ≈100, so only a thin ring of the membrane under the condensate is bent. Even a size difference too small for microscope’s resolution (≈250 nm), may then be enough for the condensates to find a favorable membrane configuration while completely overlapping. This mechanism also explains why most small condensates in the experimental images sit near the edge of the large one while fully overlapping. The ratio *R/ℓ*_*b*_ can thus be viewed as the dimensionless number to preserve when relating simulation and experiment, although the experimental value of 100 is out of reach for the Cooke model: the lipids are stable only up to Σ ≈0.5*k*_*B*_*T/σ* ^2^ [38], which would require *R* ≈600*σ*, one order of magnitude larger than the condensates we simulate and beyond what PMF calculations allow. Thus, we cannot simulate a condensate which has a *R/ℓ*_*b*_ ≫ 10 and show explicitly that the curvature coupling occurs at the rim.

### Condensate’s impact on membrane entropy

Quantifying the entropy of the membrane is computationally challenging even within the highly coarse-grained framework we use, and the change in entropy induced by a wetting condensate is likewise inaccessible with available methods. Consequently, a thermodynamically consistent value for the fluctuation-induced interaction is difficult to obtain, particularly because it is coupled to the curvature-induced interaction. What we can do instead is compare the fluctuations of the membrane’s degrees of freedom when it is bare, when it is wetted by a single condensate on one side, and when it is wetted by two condensates on opposite sides.

A lipid membrane possesses many fluctuation modes that contribute to its total entropy, ranging from covalent-bond fluctuations at small length scales to collective bending undulations at large ones. The Cooke model is highly coarsegrained—a Cooke lipid comprises only 3 beads, whereas a real lipid typically has more than ∼ 100 atoms—so fluctuations of sub-molecular degrees of freedom must be interpreted with care. The configurational space of a Cooke lipid reduces to three dimensions (one H–T–T angle and the two bond lengths H–T and T–T fully specify a configuration), far smaller than that of a real lipid. Moreover, the different modes are not truly independent but coupled, so they cannot simply be summed into a total entropy value.

The Cooke model does, however, allow us to sample long-wavelength bending undulations, and we focus on the condensate’s effect on these modes. We prepared three systems— a *bare* membrane, a membrane with a condensate on a *single side*, and a membrane with condensates on *both sides*— shown in Fig. 6(a–c). Fully covering the membrane, rather than using a condensate patch, removes the effect of lateral translation of the condensates (relative to the membrane and to each other) on the bending undulations. All simulations share the same lateral tension, Σ = 0.4 *k*_*B*_*T/σ* ^2^, the appropriate ensemble for modeling different regions of a single large suspended membrane. For each system we performed 4 independent simulations, each equilibrated for 2500 *τ*; the membrane beads were then sampled every 5 *τ*, yielding 1500 snapshots over 7500 *τ* per run. The box sizes are *L*_*xy*_ = 194.6 ± 0.2*σ*, 193.8 ± 0.2*σ*, 193.1 ± 0.2*σ* for bare, one-sided and two-sided cases, respectively. *N*_*chain*_ is set by the lipid-to-polymer ratios within the bulk of equilibrated droplets under the same conditions obtained in separate simulations. Each sampled configuration is converted to a height profile *h*^(*t*)^(*x, y*) on a uniform grid (see SI) and expressed in Fourier space as a function of wavevector,

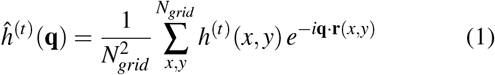

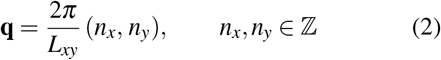

The distribution of the Fourier coefficients sampled at a given wavevector can be thought as an *entropy proxy* of that fluctuation mode (see SI). We split *ĥ*^(*t*)^(**q**) into its real and imaginary parts and, exploiting the radial isotropy of a flat membrane, bin the samples by wavevector magnitude *q*. Ideally the coefficients belonging to a mode’s exact wavevector *q*_*k*_ would be pooled together; however, because the box dimension *L*_*xy*_ fluctuates during the simulations, we instead pool all coefficients whose wavevectors lie in the immediate vicinity of *q*_*k*_, eq. (5), giving the collection *C*_*k*_ that characterizes the mode at *q*_*k*_, eq. (6).

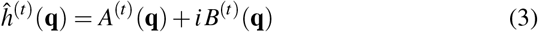

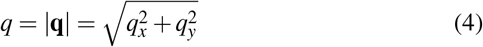

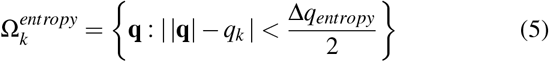

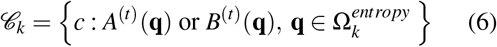

Figure 6(e) shows *C*_*k*_ at *q*_*k*_ = 0.41 *σ* ^−1^ for the three membranes; the distributions are nearly perfectly Gaussian, ensuring Δ*q*_*entropy*_ = 0.02 *σ* ^−1^ is small enough. Although they almost overlap, the two-condensate membrane has the tallest peak and the bare membrane the shortest, implying—since these are probability densities—that the bare-membrane distribution is the widest, with the most weight in its tails. This is seen more clearly in Fig. 6(f), where the bare membrane is taken as the reference and the differences from the wetted cases are plotted. The narrower distributions of the wetted membranes indicate a smaller entropy proxy for the corresponding bending-undulation mode. The discrete entropy, eq. (7), is 2.941 ± 0.003, 2.917 ± 0.004, and 2.877 ± 0.002 nats for the bare, one-condensate, and two-condensate membranes, respectively (mean ± SEM over 4 independent runs).

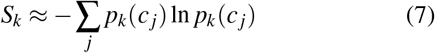

We can conclude that, overall, a wetting condensate can suppress the membrane’s bending undulation modes and the entropy associated with them. We stress, however, that the comparison is only qualitative, so whether this reduction carries a genuine thermodynamic effect remains unanswered. For example, the numerically computed discrete entropy depends strongly on the bin size used to discretize the distribution.

Because *ĥ*^(*t*)^(**q**) is already available, we can also compare the effective bending modulus of the three membranes using Helfrich’s theory [58, 59]. The mean power amplitudes |*h*(*q*_*k*_)|^2^ were computed from eqs. (8) and (9), with a bin width Δ*q*_*fit*_ = 0.03 *σ* ^−1^,

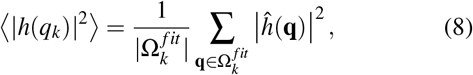

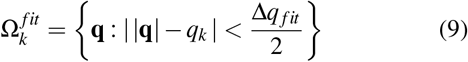

The four values from the independent runs are averaged and plotted in Fig. 6(d). According to Helfrich’s theory,

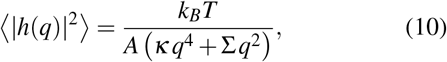

where *A* is the projected membrane area and *κ* and Σ are the effective bending modulus and lateral tension. Although the three curves nearly coincide, the bare-membrane amplitudes are the largest and the two-condensate amplitudes the smallest at most wavevectors, which is in agreement with the entropy comparison above. Fitting to eq. (10) yields the bending moduli: *κ*_0_ = 19.28 ± 0.09, *κ*_1_ = 21.31 ± 0.11, and *κ*_2_ = 23.49 ± 0.47 *k*_*B*_*T* for the bare, one-condensate, and two-condensate membranes, respectively (mean ± SEM over 4 independent runs). These are close to previously reported values for Cooke membranes and within the range for biological membranes [60, 61]. While the absolute bending moduli values obtained with this approach can be sensitive to the *hyperparameters* of the analysis, the comparison of *κ*_2_ *> κ*_1_ *> κ*_0_ is more robust as shown in Table S2.

A membrane patch wetted by a condensate can be thought of as a lipid domain with a larger effective bending modulus, so the argument of Haataja [17] for the coupling of rigid domains applies to these wetted domains as well. In their picture the registered state consists of one doubly-stiffened domain and one bare domain, and the unregistered state of two singly-stiffened domains; mode counting then gives a free-energy gain per unit area of overlap. Because we measure all three moduli directly, we can write it as

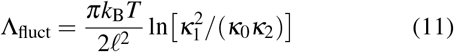

where ℓ is comparable to the bilayer thickness (≈5 nm). Registration is thus favorable only if 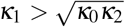, i.e. if the first condensate stiffens the membrane more than the second. Our fitted values satisfy this (*κ*_1_ = 21.31 *k*_*B*_*T* versus 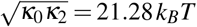), giving Λ_fluct_ ≈ 1.7 × 10^−4^ *k*_*B*_*T/*nm^2^ which would be significant for micron sized condensates. However, both the statistical and systematic uncertainties (see Table S2) are too large to determine the sign of the interaction or reliably estimate its magnitude. We therefore regard the existence and magnitude of the fluctuation-induced contribution — at both simulated and experimental sizes — as unresolved, and we do not use it to draw conclusions at either scale.

Establishing whether suppression of bending undulations can drive coupling requires separating *κ*_1_ from 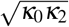 by more than the error of the fits. Varying the condensate thickness through the chain length, the polymer–lipid adhesion (*ε*_*pH*_), or the condensate density (*ε*_*AB*_) would all change the added stiffness and may open that gap. The size dependence also offers a route: Λ_fluct_ scales with the overlap area, whereas curvature-induced interactions are either size-independent — as Van Der Wel *et al*. [24] measured, and as expected from the scale invariance of the Helfrich Hamiltonian at zero tension — or, under tension, scale with the length of the curved rim. A crossover radius beyond which fluctuations dominate must therefore exist, but locating it would require a value of Λ_fluct_ we do not have.

What our data do support is the curvature-induced mechanism. The PMF behavior and observed membrane geometries are consistent with curvature-dominated coupling. It accounts for the experimental observation that the smaller condensate sits near the rim of the larger one, which the fluctuation mechanism — depending only on overlap area — cannot explain.

### Model limitations

There is an extensive body of theory and closed-form expressions for membrane-mediated interactions [12, 14, 62– 65], but most of it does not apply cleanly to our system. These expressions are typically derived in the small-deformation regime (a linearized approximation of the full Helfrich energy), often neglect tension, and are valid only at large separations relative to the inclusion size (*r*_*xy*_ ≫ *R*). The last assumption is the one that breaks most severely here: we are interested in the short-range regime *R* ∼ *r*_*xy*_, and our condensates can overlap completely (*r*_*xy*_ →0), so they cannot be treated as point-like curvature sources [66]. Our curvature-inducing object is also not rigid–it responds to stress–and the two interacting condensates can have a size asymmetry. The shape of the condensates can deviate from a perfect disk due to thermal fluctuations or as a response to the curvature induced by the other condensate. It is for all of these reasons that we turn to numerical simulations.

Our model is generic, and the chemistry of the protein– lipid interface only enters it through a few effective parameters. Lipids and proteins can interact in many specific ways [67, 68], and how strongly a condensate wets a membrane depends on the chemical details of both [69], on the IDP sequence [70, 71], and on solvent conditions such as pH and ion concentration. The same factors also set the condensate– solvent interfacial tension Σ_*cs*_, and therefore the curvature that a condensate creates on the membrane. Since our analysis is written in terms of *κ*, Σ and *R*, this chemistry is not really lost: sequence, lipid composition and solvent conditions can be thought of as setting these parameters. Two simplifications should still be pointed out. First, following the experiments that use His-tags [9], our chains are tethered to the lipid head beads, so the wetting is chemical rather than physical (generic, non-bonded) adhesion. This kind of anchoring is also used in nature [72], and some of our findings should still apply, but care is needed when extending them to purely physical wetting. Second, our membrane is homogeneous, while a real membrane contains different lipid domains and embedded proteins, which also affect where and whether condensates form [73]. We restrict ourselves to a chemically uniform membrane and do not alter its characteristics by varying the coarse-grained interaction parameters of the lipid particles.

Our condensates are also much denser than real ones. Since we model them as homopolymers with generic attractions, they form liquid but compact droplets with a mesh size close to the monomer size. Real IDP condensates are semi-dilute and much more permeable; LAF1-RGG droplets, for example, have a mesh size of 3 − 8 nm [74]. A looser condensate has might have lower Σ_*cs*_ and possibly makes the membrane patch it wets less stiff, so it might create less curvature and also suppress fewer bending modes. The interactions we report are therefore closer to an upper bound for what a condensate of a given size can do to the membrane, for both mechanisms. Getting such loose and permeable droplets requires the more specific and localized interactions of residue-level coarse-grained models with several bead types [36]. These models come with many more parameters, they are harder to couple to a generic lipid model, and they still miss the steric effects of explicit side chains.

The last group of limitations comes from the low resolution of our lipid model. Coarse-graining removes some degrees of freedom. The entropy of the removed modes ends up inside the effective interactions between the beads, where it shows up as energy instead of entropy. This matters most for the entropy–energy competition that is central to this work, with Cooke’s model we cannot draw conclusions about the molecular-scale modes that it cannot resolve. The bending undulations we analyzed above are collective enough to survive the coarse-graining, but a similar comparison of molecular-level configurational entropies [75] would need a lipid model with more detail.

## IV. CONCLUSIONS

In this work we used coarse-grained molecular dynamics simulations to explain the transmembrane coupling of protein condensates seen in recent experiments [9, 10]. In those experiments, micron-sized condensates on opposite surfaces of a suspended flat lipid membrane prefer to sit fully overlapped, even though they cannot touch each other. This points to an indirect, membrane-mediated interaction between them. There are two candidate mechanisms: a curvature-induced interaction, which is energetic in origin, and a fluctuation-induced interaction, which is entropic. To probe them, we combined Cooke’s implicit-solvent lipid model with a generic bead-spring polymer model for the condensate.

We first computed the potential of mean force between two condensates of the same size. Fully overlapping configurations turned out to be unfavorable. Instead the condensates prefer to sit partially overlapped, bending the membrane into an S-like shape, which agrees with previous studies. When we increased the size ratio between the two condensates, full overlap became more favorable. We propose the following explanation. Because wetting is driven by the tethers, the condensate spreads into a thin film on the membrane, so it exerts no significant bulk pressure and imposes curvature only along its rim, where it meets the solvent. Membrane tension then damps this deformation away from the rim. The result is a flat membrane region under the middle of the condensate, ringed by a curved rim. A smaller condensate on the opposite side of the membrane can be trapped in that ring if it adheres to it.

Quantifying the entropic contribution is harder. We compared the entropy of the bending-undulation modes of three membranes: a bare one, one wetted by a single condensate, and one wetted by condensates on both sides. We also estimated an effective bending modulus for each. Both the first and the second condensate suppress the undulation modes and stiffen the membrane. Whether the overlap of two condensates is entropically favorable, however, remains inconclusive.

Overall, we conclude that full overlap of the two condensates can be thermodynamically favorable given the right combination of wetting mechanism and membrane tension. This hypothesis should be tested against experiments and further simulations. Continuum methods in particular would be valuable for understanding the curvature-induced mechanism and how it depends on the basic parameters of the system: the bending modulus of the membrane, the mechanical lateral tension, the condensate-solvent interfacial tension, and the condensate size.

## ACKNOWLEDGEMENTS

The authors acknowledge support from the National Science Foundation under Grant No. BIO-2529782, the National Institutes of Health under Grant No. R35GM139531 (J.S.), and the Welch Foundation under Grant No. F-2257 (J.S.).

## SUPPLEMENTAL INFORMATION

The supplementary information provides details on potential of mean force calculation and bending undulation analysis.

