## Supplementary material for "Transmembrane coupling of protein condensates via membrane-mediated interactions: A simulation study": Supplmental Information

### CONTENTS

I. Potential of the Mean Force Calculations 2

II. Bending Fluctuation Analysis 3

III. Scale-invariance and bending screening with tension 7

References 7

*

### I. POTENTIAL OF THE MEAN FORCE CALCULATIONS

**Reaction coordinate:** We chose the in-plane center-of-mass distance between the two condensates as the reaction coordinate,

$$r_{xy} \;=\; \left\|\mathbf{R}_{xy}^{\text{upper}} - \mathbf{R}_{xy}^{\text{lower}}\right\|, \tag{S1}$$

where $\mathbf{R}_{xy}^{\text{upper}}$ and $\mathbf{R}_{xy}^{\text{lower}}$ are the projections onto the membrane plane $(xy)$ of the centers of mass of the polymer chains tethered to the upper and lower leaflets, respectively. The vertical component $(z)$ is excluded so that the bias acts only along the membrane plane.

**Biasing potential and choice of windows:** To sample values of $r_{xy}$ that would be rare in an unbiased run, we used umbrella sampling. In each window, an extra harmonic potential is added to the system,

$$U_{\text{bias}}(r_{xy}, r_0) \;=\; \tfrac{1}{2}\, k_{\text{umb}}\, (r_{xy} - r_0)^2, \tag{S2}$$

which keeps the condensate–condensate distance close to a chosen center $r_0$ while still letting it fluctuate. We used a spring constant of $k_{\text{umb}} = 25\, k_B T/\sigma^2$, where $\sigma$ is the diameter of a lipid tail bead as usual. For two condensates of 500 polymer chains, we placed window centers from $r_0 = 0.0\,\sigma$ to $r_0 = 65.0\,\sigma$ in steps of $0.5\,\sigma$. Different ranges of $r_0$ have been used for different systems. The spacing was chosen small enough that the histograms of $r_{xy}$ from neighboring windows overlap, which is required for the reweighting step.

**Per-window simulation protocol :** Each window was simulated as part of the same pipeline used for the unbiased system: first the membrane was relaxed to target temperature and tension, then the polymers were tethered and relaxed, and finally the production run was performed in the $N\Sigma T$ ensemble with the lateral pressure set by the target tension. The bias in Eq. (S2) was switched on at the end of the polymer-relaxation stage and kept on for the rest of the run. The simulation time step was $\Delta t = 0.005\,\tau$.

After switching the bias on, each window was first equilibrated for $5 \times 10^5$ steps, and then a production run was performed during which the value of $r_{xy}$ was written to disk every 100 steps. The total production length was typically around $5 - 50 \times 10^6$ steps per window, depending on the correlation time of $r_{xy}$ at that window, as explained later. For each window, several independent replicas were run with different random seeds; the time series of $r_{xy}$ from all replicas at the same $r_0$ were concatenated before analysis.

**Free-energy reconstruction with WHAM :** The biased histograms from all windows were combined into a single unbiased free-energy profile $F(r_{xy})$ using the weighted histogram analysis method (WHAM) [1, 2]. We used the implementation of Grossfield (`wham`, version 2.1.0) [3], run in reduced (LJ) units. The correlation time $\tau_{correl}$ was calculated from the time series of $r_{xy}$, in order to perform the error analysis (see below).

WHAM finds the set of free-energy offsets $\{F_i\}$ between windows that makes the windows mutually consistent, and then returns the unbiased probability distribution $P(r_{xy})$ on a chosen grid. We used 300 bins covering the full sampled range of $r_{xy}$ and a convergence tolerance of $10^{-4}$ on the $F_i$ values. The potential of the mean force is obtained from

$$F_{\text{PMF}}(r_{xy}) = -k_B T \ln P(r_{xy}) + k_B T \ln r_{xy}, \tag{S3}$$

and is reported up to an additive constant (we set the minimum of $F_{\text{PMF}}(r_{xy})$ to zero). The term $k_B T \ln r_{xy}$ is the Jacobian of the chosen reaction coordinate.

**Statistical errors via correlation times and Monte Carlo bootstrap :** The samples in the time series of $r_{xy}$ within a window are not statistically independent: consecutive frames are correlated, so the true number of independent measurements is smaller than the raw sample count. To take this into account we estimated the integrated autocorrelation time $\tau_{\text{int}}$ for each window and passed it to WHAM, where it is used by the bootstrap procedure to set the effective sample size.

For each window, we computed the autocorrelation function of the $r_{xy}$ time series,

$$C(t) \;=\; \frac{\langle\, \delta r_{xy}(t')\, \delta r_{xy}(t'+t) \rangle}{\langle\, \delta r_{xy}^2 \rangle}, \tag{S4}$$

where $\delta r_{xy}(t) = r_{xy}(t) - \langle r_{xy} \rangle$ and the brackets denote a time average. The ACF was evaluated through an FFT on the mean-subtracted series. The integrated autocorrelation time was then obtained as

$$\tau_{\text{int}} \;=\; \tfrac{1}{2} \;+\; \sum_{t=1}^{M} C(t), \tag{S5}$$

with the cutoff $M$ chosen using Sokal's automatic windowing procedure [4]: $M$ is the smallest lag for which $M \geq c\,\tau_{\rm int}(M)$. We tried several values of $c$ $(6, 8, 9, 10)$ and confirmed that M converges around $c = 9, 10$.

The correlation timescale, $\tau_{correl} = 2\tau_{int}$, depends strongly on the window, varying by more than an order of magnitude across different $r_{xy}$ values. For example, to obtain 100 independent samples at a window where $\tau_{int} \approx 1500$, the simulation must run for $1500 \times 2 \times 100 \times 100 = 3 \times 10^7$ timesteps, the second 100 factor is included since $r_{xy}$ is recorded every 100 timesteps. $\tau_{correl}$ is more commonly called the statistical inefficiency, $g$ [5]. The effective number of independent samples per window was then estimated as

$$N_{\rm eff} = \frac{N}{2\,\tau_{\rm int}}, \tag{S6}$$

where $N$ is the number of stored frames in the window. Windows with multiple replicas at the same $r_0$ were treated as a single longer time series for this calculation. We checked that all windows had $N_{\rm eff} \gtrsim 100$, which is the minimum we considered acceptable for the WHAM input.

The statistical uncertainty on $F_{\rm PMF}(r_{xy})$ was then obtained with the Monte Carlo bootstrap procedure built into the WHAM code [3]. For each bootstrap trial, a new synthetic histogram is generated for every window by drawing $N/\tau_{\rm int}$ random numbers from the cumulative distribution defined by that window's actual histogram (the division by $\tau_{\rm int}$ accounts for correlations and prevents the bootstrap from underestimating the error). WHAM is then run on the full set of synthetic histograms to produce one "bootstrap" free energy profile. We set the *num_ MC_ trials* parameter to 300 and took the standard deviation of the resulting $F_{\rm PMF}(r_{xy})$ values, bin by bin, as our estimate of the statistical uncertainty.

### II. BENDING FLUCTUATION ANALYSIS

The effective bending modulus of the bilayer was extracted from the equilibrium spectrum of out-of-plane height fluctuations, computed from the production $N\Sigma T$ trajectory described in methods section. The pipeline has three stages: (i) per-frame extraction of a smooth midplane height field $h(\mathbf{r}(x, y), t)$ from particle coordinates; (ii) computation of the time-averaged Fourier spectrum $\langle|\hat{h}(\mathbf{q})|^2\rangle$ on a wave-vector grid; and (iii) fit of the spectrum to the Helfrich expression to recover an effective bending modulus $\kappa_{eff}$. From the same Fourier spectrum, we additionally compute a quasi per-mode differential-entropy proxy.

We were conservative in choosing the wave-number range. Since our goal is not to obtain a single $\kappa_{eff}$ value but rather statistically meaningful power spectra for three different systems (bare membrane, membrane with one condensate, and membrane with two condensates), we picked $q^{\rm min}$ for the Helfrich fit by excluding lower $q$ values where the mean power amplitudes fell within each other's error bars. The default values used for analysis are given in Table S1.

Table S1: Default parameters used in the bending fluctuation analysis.

| Symbol | Meaning | Default value |
|---|---|---|
| $\lambda^{\rm max}$ / $q^{\rm min}$ | lower bound for wave-number for Helfrich fit | $78.5\,\sigma$ / $0.08\,\sigma^{-1}$ |
| $\lambda^{\rm min}$ / $q^{\rm max}$ | upper bound for wave-number for Helfrich fit | $19.5\,\sigma$ / $0.32\,\sigma^{-1}$ |
| $\Delta q_{fit}$ | bin width of $q$ for Helfrich fit | $0.03\,\sigma^{-1}$ |
| $\sigma_g$ | width of gaussian filter for height grid calculation | $0.8\sigma$ |
| $a$ | real-space grid spacing of $h(\mathbf{r}_\parallel)$ | $0.5\,\sigma$ |
| $u_z^{\rm thr}$ | threshold for assigning lipids to a leaflet | $0.4\,\sigma$ |

**Per-frame mid-plane height field:** For each saved frame we construct a smooth periodic height field $h^{(t)}(\mathbf{r}(x, y))$, defined on a regular grid that covers the simulation cell. To assign lipid chains to leaflets we first compute the unit vector $\hat{u} = (u_x, u_y, u_z)$ pointing from the midpoint of the two tail beads to its head bead. A lipid is assigned to the upper leaflet if $\hat{u}_z > u_z^{\rm thr}$ and to the lower leaflet if $\hat{u}_z < -u_z^{\rm thr}$; lipids with $|\hat{u}_z| \leq u_z^{\rm thr}$ are deemed ambiguous (severely tilted, mid-flip, or transiently dissociated) and excluded from the height field for that frame. The parameter $u_z^{\rm thr}$ is intentionally generous: at the operating temperature $k_{\rm B}T = \varepsilon$ the equilibrium tilt distribution is narrow and only a small fraction of lipids $(0.3 - 0.5\%)$ are excluded at each frame.

Lipids that have detached from the bilayer would contaminate the height field with spurious z-values, so these chains must be excluded from the membrane's height profile calculations. This is not a problem for most lipid models, but the coarse representation of Cooke's lipid chains lets them leave the membrane through thermal fluctuations. For each leaflet separately, we therefore build a periodic cKDTree on the candidate head positions and retain only the largest connected component of the graph in which two heads are linked when their inter-head distance is below a cluster cutoff $r_{\rm clu} = 1.3\sigma$. This cutoff is roughly the first minimum of the lipid–lipid head–head pair-correlation function in the assembled bilayer; it is comfortably larger than the WCA contact distance and small enough to disconnect any lipid whose head has moved more than a single shell out of the bilayer plane. The clustering step removes roughly $0.8 - 1.3\%$ of lipids from each leaflet.

After each lipid chain is assigned to a leaflet, a smooth height field is obtained from the head positions as a density-normalized kernel-weighted average of the head $z$-coordinates. The construction is somewhat inspired by particle–mesh methods, in which particle data are deposited onto a regular grid, and follows a square-mesh approach similar to that of [6]. Let $\{\mathbf{r}_i^{(\alpha)}\}$ denote the retained head positions in leaflet $\alpha \in \{\text{upper}, \text{lower}\}$, with components $(x_i, y_i, z_i)$. The simulation cell is partitioned into an $N_{\rm grid} \times N_{\rm grid}$ grid with $N_{\rm grid} = \text{round}(L_{xy}/a)$ and spacing $a = 0.5\sigma$ (so $N_{\rm grid} \approx 400$ for the production runs at $L_{xy} \approx 200\sigma$; in practice $a$ fluctuates within $0.5 \pm 0.0006\,\sigma$ and $N_{\rm grid}$ within $388 \pm 2$). The leaflet height field is then

$$h^{(\alpha)}(\mathbf{r}_{xy}) = \frac{\sum_{i\in\alpha} z_i\, K\Big(\mathbf{r}_{xy} - \mathbf{r}_{xy,i}^{(\alpha)}\Big)}{\sum_{i\in\alpha} K\Big(\mathbf{r}_{xy} - \mathbf{r}_{xy,i}^{(\alpha)}\Big)},$$

with denominator always non-zero in our analysis. The weighting kernel $K = G_{\sigma_g} * W$ is a bilinear (tent) assignment kernel $W$ smoothed by an isotropic periodic Gaussian $G_{\sigma_g}$,

$$W(\mathbf{r}(x,y)) = \Big(1 - \tfrac{|x|}{a}\Big)_+ \Big(1 - \tfrac{|y|}{a}\Big)_+, \qquad G_{\sigma_g}(\mathbf{r}(x,y)) = \frac{1}{2\pi\sigma_g^2}\exp\left(-\frac{|\mathbf{r}(x,y)|^2}{2\sigma_g^2}\right),$$

where $(u)_+ \equiv \max(u, 0)$ and $a$ is the grid spacing. The Gaussian has width $\sigma_g = 0.8\sigma$, truncated at $r_g = 1.5\sigma$.

The midplane field is taken as the arithmetic mean of the two leaflet fields,

$$h(\mathbf{r}_{xy}, t) = \tfrac{1}{2}\big[h^{\rm upper}(\mathbf{r}_{xy}, t) + h^{\rm lower}(\mathbf{r}_{xy}, t)\big].$$

Per frame we save $h(\cdot, t)$, the grid coordinate vector, and the instantaneous lateral box length $L_{xy}(t)$. The grid and box size must be tracked, because the barostat allows $L_{xy}$ to fluctuate; failing to do so would shift the q-grid relative to the data and broaden the binned spectrum. The analysis based on power spectrum of simulated membranes is notoriously sensitive to how the analysis is performed [7]. In Table S2, we show that while $\kappa$ values can be different, the trend is the same except when $q$ bin width is $0.04\sigma^{-1}$.

**Per-mode entropy proxy calculation :** In this section we explain how we perform an approximate comparison of the bending undulation *entropy proxy* of the free membranes and membranes with condensates.

The entropy of a membrane height field (h(x,y)) should be understood as the entropy of the probability distribution over entire height-field configurations, not merely the entropy of the distribution of local height values. In real space, a gridded height field is represented by many coupled variables, one for each grid point. Therefore, the direct entropy calculation would require estimating the full joint probability distribution of all height values on the grid. Even for a finite grid, this is a very high-dimensional density-estimation problem, and it rapidly becomes impractical. This is the main reason that a direct real-space calculation of the configurational entropy of the height field is extremely difficult, even for numerical, non-thermodynamic entropy.

A Fourier transform provides a more convenient basis for this problem. For each sampled height field, the transformation from (h(x,y)) to its Fourier coefficients $(h_{\mathbf{q}})$ is an exact change of coordinates within the chosen resolution and cutoff range. Thus, no information is lost by working in Fourier space. However, the exactness of the Fourier decomposition should not be confused with statistical independence. Fourier modes are orthogonal basis functions, but the corresponding random coefficients can still be statistically correlated in the ensemble of thermally fluctuating membranes.

In general, the full membrane free-energy functional can contain nonlinear geometric terms. When such terms are expressed in Fourier space, they generate interactions between different wavevectors. In that case, the probability distribution of the height field does not factorize into independent distributions for each $(h_{\mathbf{q}})$, and the total entropy cannot be obtained exactly by summing single-mode entropies. The sum of single-mode entropies is exact only when the joint distribution over Fourier coefficients factorizes into a product over modes.

The usual independent-mode picture arises from the small-gradient approximation to the Helfrich Hamiltonian as explained in [8]. In this approximation, the membrane energy becomes quadratic in the height field and diagonal in Fourier space. The Hamiltonian can then be written as a sum of separate contributions from different wavevectors. This removes the physical/statistical coupling between modes at the level of the quadratic theory. As a result, the Boltzmann probability distribution factorizes over Fourier modes.

For a quadratic membrane Hamiltonian of the form $\frac{A}{2}\sum_{\mathbf{q}}\left(\kappa q^4+\Sigma q^2\right)|h_{\mathbf{q}}|^2$, where A is the projected membrane area, the equilibrium probability is $\frac{1}{Z}\exp[-\beta H[h]]$. Writing a complex Fourier coefficient as $h_{\mathbf{q}} = a_{\mathbf{q}} + ib_{\mathbf{q}}$, we have $|h_{\mathbf{q}}|^2 = a_{\mathbf{q}}^2 + b_{\mathbf{q}}^2$. Therefore, for each independent nonzero Fourier mode, $P_{\mathbf{q}}(a_{\mathbf{q}}, b_{\mathbf{q}}) \propto \exp\left[-\frac{\beta A}{2}\left(\kappa q^4+\Sigma q^2\right)\left(a_{\mathbf{q}}^2+b_{\mathbf{q}}^2\right)\right]$. This is a two-dimensional Gaussian distribution in the real and imaginary parts of $(h_{\mathbf{q}})$. Thus, within the quadratic small-gradient Helfrich approximation, $(a_{\mathbf{q}})$ and $(b_{\mathbf{q}})$ are Gaussian-distributed variables, and different Fourier modes are statistically independent.

This also clarifies the distinction between the Fourier coefficient distribution and the fluctuation spectrum. The quantities $(a_{\mathbf{q}})$ and $(b_{\mathbf{q}})$ are Gaussian in the quadratic theory, but the power $(|h_{\mathbf{q}}|^2 = a_{\mathbf{q}}^2 + b_{\mathbf{q}}^2)$ is not Gaussian. The power spectrum used in Helfrich fits is based on averages of $(|h_{\mathbf{q}}|^2)$, whereas an entropy-proxy calculation should be based on the distribution of the Fourier coefficients themselves.

Therefore, the distribution of Fourier coefficients at a given wavevector, or within a narrow shell of wavevectors with the same magnitude $(q = |\mathbf{q}|)$, can be used to estimate the entropy associated with that mode or group of modes. If the independent-mode approximation is valid, the entropy of the height field over a chosen range of wavelengths can be approximated as the sum of the entropies of the retained Fourier modes. If mode coupling is significant, this sum overestimates the true entropy, because the joint entropy is reduced by statistical correlations between modes.

In Fig. S1, we show that the distribution of $c_q = a_q \cup b_q$ can be very well fitted with a gaussian of zero mean.

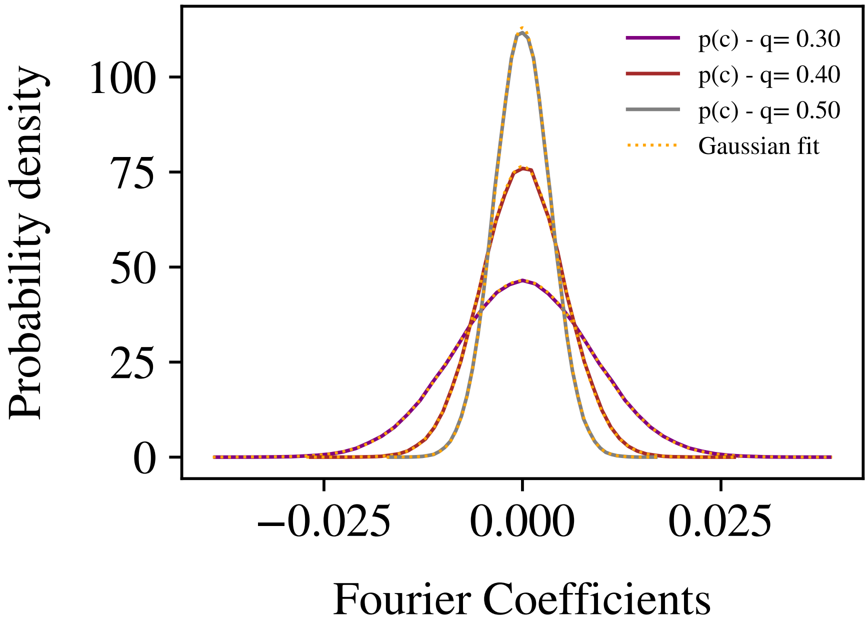


Figure S1: The distribution of fourier coefficients of the sampled height field $\hat{h}^{(t)}(\mathbf{q}) = A^{(t)}(\mathbf{q}) + i\,B^{(t)}(\mathbf{q})$ at different wavevectors $q = |\mathbf{q}| = \{0.3, 0.4, 0.5\}\sigma^{-1}$ are plotted. The distribution can be fitted very well with a gaussian centered at zero, in agreement with the theory. $\Delta q$ is $0.02\sigma^{-1}$, bin centers are given in the legend. Bare membrane.

**Fitting for bending modulus :** As mentioned previously, there are some arbitrary 'hyperparameter' choices that we have made for the entropy and bending undulation analysis. In Table S2 we show that while absolute values for fitted $\kappa$ are affected by the arbitrary choices, the $\kappa_2 > \kappa_1 > \kappa_0$ trend is robust except at $\Delta q = 0.04\sigma^{-1}$. The error estimation is done using fit results from 4 independent runs. Both statistical (among 4 independent runs) and systematical (across different analysis parameters) are too large to estimate a log ratio.

Table S2: Robustness of the fitted bending modulus $\kappa$ and effective tension $\Sigma$ to the choice of hyperparameters using $[0.08, 0.32]\sigma^{-1}$ range for $q$. Each row denotes the parameter(s) changed from the default value (top row), with all others held at their defaults. Subscripts denote the bare membrane (0), one condensate (1) and two condensates (2); While the absolute values of the bending moduli are sensitive to the parameter choices, the relative trend of $\kappa_2 > \kappa_1 > \kappa_0$ is more robust. Uncertainties are 1 standard deviation SEM of four independent runs fitted separately; the uncertainty on the log ratio is obtained by standard error propagation, $\sigma_R = \sqrt{(2\sigma_{\kappa_1}/\kappa_1)^2 + (\sigma_{\kappa_2}/\kappa_2)^2 + (\sigma_{\kappa_0}/\kappa_0)^2}$.

| | $\kappa$ | | | $\Sigma$ | | | |
|---|---|---|---|---|---|---|---|
| Configuration | $\kappa_0$ | $\kappa_1$ | $\kappa_2$ | $\Sigma_0$ | $\Sigma_1$ | $\Sigma_2$ | $\ln(\kappa_1^2/(\kappa_0\kappa_2))$ |
| Default | $19.28 \pm 0.09$ | $21.31 \pm 0.11$ | $23.49 \pm 0.47$ | $0.348 \pm 0.006$ | $0.318 \pm 0.004$ | $0.304 \pm 0.020$ | $0.003 \pm 0.023$ |
| $a = 0.35$ | $19.11 \pm 0.09$ | $21.14 \pm 0.12$ | $23.33 \pm 0.47$ | $0.350 \pm 0.006$ | $0.320 \pm 0.004$ | $0.305 \pm 0.020$ | $0.003 \pm 0.023$ |
| $a = 0.8$ | $19.52 \pm 0.09$ | $21.57 \pm 0.11$ | $23.75 \pm 0.47$ | $0.346 \pm 0.006$ | $0.316 \pm 0.004$ | $0.301 \pm 0.020$ | $0.003 \pm 0.023$ |
| $u_z^{thr} = 0.5,\ a = 0.8$ | $19.61 \pm 0.09$ | $21.66 \pm 0.11$ | $23.85 \pm 0.47$ | $0.345 \pm 0.006$ | $0.315 \pm 0.004$ | $0.301 \pm 0.020$ | $0.003 \pm 0.023$ |
| $u_z^{thr} = 0.6,\ a = 0.8$ | $19.76 \pm 0.09$ | $21.81 \pm 0.11$ | $24.00 \pm 0.47$ | $0.344 \pm 0.006$ | $0.314 \pm 0.004$ | $0.300 \pm 0.020$ | $0.003 \pm 0.023$ |
| $\sigma_g = 0.6,\ a = 0.8$ | $19.06 \pm 0.09$ | $21.08 \pm 0.11$ | $23.24 \pm 0.47$ | $0.350 \pm 0.006$ | $0.320 \pm 0.005$ | $0.306 \pm 0.020$ | $0.003 \pm 0.023$ |
| $\sigma_g = 1.2,\ a = 0.8$ | $20.17 \pm 0.09$ | $22.26 \pm 0.11$ | $24.49 \pm 0.47$ | $0.340 \pm 0.006$ | $0.310 \pm 0.004$ | $0.295 \pm 0.019$ | $0.003 \pm 0.022$ |
| $\Delta q = 0.02$ | $16.77 \pm 0.12$ | $18.60 \pm 0.07$ | $20.24 \pm 0.36$ | $0.463 \pm 0.009$ | $0.421 \pm 0.008$ | $0.431 \pm 0.022$ | $0.019 \pm 0.021$ |
| $\Delta q = 0.04$ | $16.66 \pm 0.38$ | $19.28 \pm 0.54$ | $18.89 \pm 1.26$ | $0.449 \pm 0.012$ | $0.405 \pm 0.016$ | $0.444 \pm 0.039$ | $0.166 \pm 0.090$ |

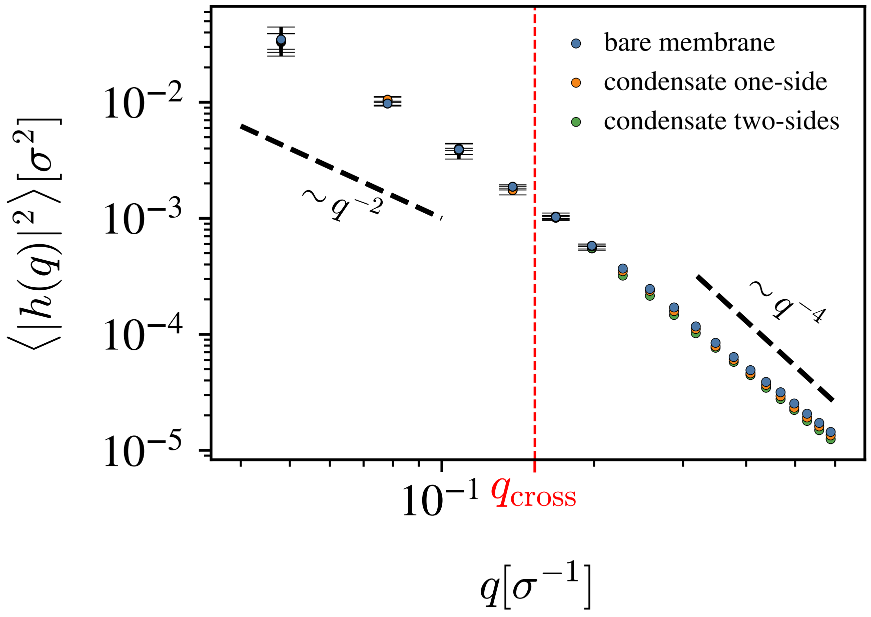


Figure S2: The full fluctuation spectrum data that high $q$ values where continuum approximation starts breaking. The gradual switch from bending ($\sim q^{-4}$) dominated to tension dominated ($\sim q^{-2}$) regime as $q$ gets smaller is visible.

### III. SCALE-INVARIANCE AND BENDING SCREENING WITH TENSION

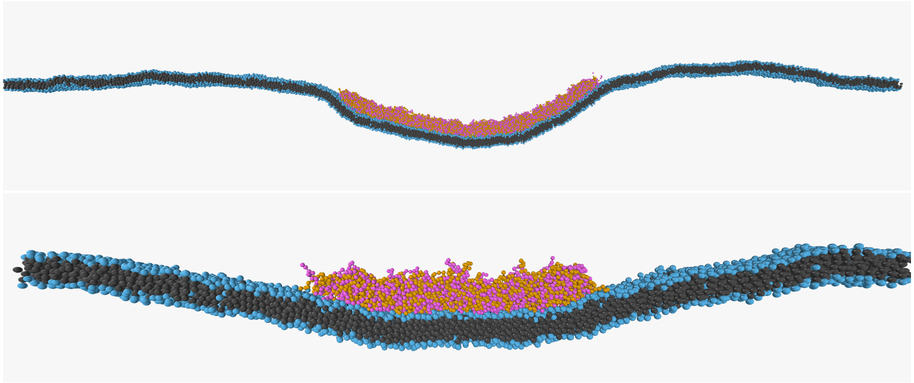

Figure S3: Two cross-sectional snapshots with tensionless ($\Sigma = 0$) membrane and a condensate. *Upper figure:* Large system with $L_{xy} \approx 450\sigma$, $N_{chain} = 4000$, $R \approx 70\sigma$. *Lower figure:* Small system with $L_{xy} \approx 150\sigma$, $N_{chain} = 444$, $R \approx 23\sigma$. The curvature imposed by the condensate deforms the whole membrane from its flat position for both systems.

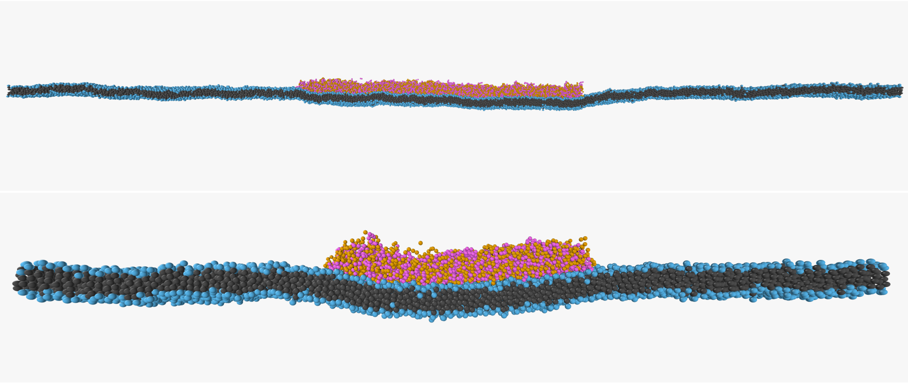

Figure S4: Two cross-sectional snapshots with $\Sigma = 0.4 k_B T/\sigma^2$ membrane and a condensate. *Upper figure:* Large system with $L_{xy} \approx 450\sigma$, $N_{chain} = 4000$, $R \approx 70\sigma$. *Lower figure:* Small system with $L_{xy} \approx 150\sigma$, $N_{chain} = 444$, $R \approx 23\sigma$. The curvature imposed by the condensate deforms a small region at the condensate rim for the larger system ($R/\ell_b \approx 10.0$) while the deformation spreads to the most of the membrane under the condensate for small system ($R/\ell_b \approx 3.3$).
